# A transcriptional cycle suited to daytime N_2_ fixation in the unicellular cyanobacterium Candidatus *Atelocyanobacterium thalassa* (UCYN-A)

**DOI:** 10.1101/469395

**Authors:** María del Carmen Muñoz-Marin, Irina N. Shilova, Tuo Shi, Hanna Farnelid, Ana Maria Cabello, Jonathan P. Zehr

**Author notes:** Address correspondence to Prof. Jonathan Zehr. Present address: Second Genome, Inc, South San Francisco, CA, 94080, USA.

## Abstract

The symbiosis between a marine alga and a N_2_-fixing cyanobacterium (UCYN-A) is geographically widespread in the oceans and is important in the marine N cycle. UCYN-A is uncultivated, and is an unusual unicellular cyanobacterium because it lacks many metabolic functions, including oxygenic photosynthesis and carbon fixation, which are typical in cyanobacteria. It is now presumed to be an obligate symbiont of haptophytes closely related to *Braarudosphaera bigelowii*. N_2_-fìxing cyanobacteria use different strategies to avoid inhibition of N_2_ fixation by the oxygen evolved in photosynthesis. Most unicellular cyanobacteria temporally separate the two incompatible activities by fixing N_2_ only at night, but surprisingly UCYN-A appears to fix N_2_ during the day. The goal of this study was to determine how the unicellular UCYN-A coordinates N_2_ fixation and general metabolism compared to other marine cyanobacteria. We found that UCYN-A has distinct daily cycles of many genes despite the fact that it lacks two of the three circadian clock genes found in most cyanobacteria. We also found that transcription patterns in UCYN-A are most similar to marine cyanobacteria that are capable of aerobic N_2_ fixation in the light such as *Trichodesmium* and heterocyst-forming cyanobacteria, rather than *Crocosphaera* or *Cyanothece* species, which are more closely related to unicellular marine cyanobacteria evolutionarily. Our findings suggest that the symbiotic interaction has resulted in a shift of transcriptional regulation to coordinate UCYN-A metabolism with the phototrophic eukaryotic host, thus allowing efficient coupling of N_2_ fixation (by the cyanobacterium) to the energy obtained from photosynthesis (by the eukaryotic unicellular alga) in the light.

**Importance:** The symbiotic N_2_-fixing cyanobacterium UCYN-A and its eukaryotic algal host, which is closely related to *Braarudosphaera bigelowii*, have been shown to be globally distributed and important in open ocean N_2_ fixation. These unique cyanobacteria have reduced metabolic capabilities, even lacking genes for oxygenic photosynthesis and carbon fixation. Cyanobacteria generally use energy from photosynthesis for nitrogen fixation, but require mechanisms for avoiding inactivation of the oxygen-sensitive nitrogenase enzyme by ambient oxygen (O_2_) or the O_2_ evolved through photosynthesis. This study shows that the symbiosis between the N_2_-fixing cyanobacterium UCYN-A and its eukaryotic algal host has led to adaptation of its daily gene expression pattern in order to enable daytime aerobic N_2_ fixation, which is likely more energetically efficient than fixing N_2_ at night, as in other unicellular marine cyanobacteria.

## Introduction

Nitrogen (N_2_)-fixing microorganisms (diazotrophs), which reduce atmospheric N_2_ to biologically available ammonium, are critical components of aquatic and terrestrial ecosystems because they supply fixed inorganic N (1). Cyanobacteria are particularly important in N_2_ fixation because they can fuel the energy intensive N_2_ reduction reaction using energy supplied by oxygenic photosynthesis. In the oceans, the filamentous, non-heterocyst-forming cyanobacterium *Trichodesmium* and the heterocyst-forming symbiont of diatoms (*Richelia* and related cyanobacteria) were believed to be the major N_2_-fixing microorganisms until the discovery of the unicellular cyanobacteria *Crocosphaera, Cyanothece* and Candidatus *Atelocyanobacterium thalassa* (UCYN-A) in the open ocean. *Crocosphaera* and *Cyanothece* are free-living marine cyanobacteria, but UCYN-A is unusual in that it lacks oxygenic photosynthesis and is a symbiont of a haptophyte alga (related to *Braarudosphaera bigelowii*). The UCYN-A symbiosis is geographically widespread and is important in oceanic N_2_ fixation (2–5). The UCYN-A genome has been greatly reduced, with massive metabolic streamlining including the loss of the oxygen-evolving Photosystem II (PSII), the carbon-fixing enzyme RuBisCO, and the entire tricarboxylic acid (TCA) cycle (6). UCYN-A has been shown to supply fixed N to the haptophyte in exchange for fixed carbon (4, 7), but it is not known how these two single-celled organisms coordinate metabolism and cell growth over the daily division cycle.

N_2_ fixation requires energy and reductant, but the nitrogenase enzyme is inactivated by oxygen (O_2_). Cyanobacteria generally have access to sufficient energy from photosynthesis but require mechanisms for avoiding inactivation of nitrogenase and N_2_ fixation by ambient oxygen (O_2_) or the O_2_ evolved through photosynthesis. *Trichodesmium* and heterocystous cyanobacteria such as *Richelia* and *Nostoc* fix N_2_ during the day, whereas the free-living unicellular *Crocosphaera* and *Cyanothece* fix N_2_ at night. Interestingly, the symbiotic UCYN-A appears to fix N_2_ during the day (8–10), in contrast to most other unicellular marine N_2_-fixing cyanobacteria, such as *Crocosphaera* and *Cyanothece*.

The processes of N_2_-fixation and photosynthesis in cyanobacteria are regulated daily to increase cellular fitness and ecological competitiveness (11–13). Most cyanobacteria have circadian rhythms (11, 14, 15) that are involved in controlling daily cycles of gene transcription and protein synthesis by signal transduction pathways involving the circadian clock *kai* genes. UCYN-A lacks two of the three *kai* genes (*kaiA* and *kaiB*) known in most other cyanobacteria, whereas the non-N_2_-fixing cyanobacterium *Prochlorococcus* only lacks *kaiA*. Thus, the daily whole genome expression pattern in UCYN-A is of interest to determine if there are daily patterns as in all other cyanobacteria compared to evolutionarily-related unicellular cyanobacteria.

We used a whole genome transcription array that targets two genetically distinct uncultivated sub-lineages of UCYN-A (UCYN-A1 and UCYN-A2), which have similar, but genetically distinct hosts. We compared the UCYN-A whole genome diel transcription patterns to those of *Cyanothece* sp. ATCC 51142 (16) and *Crocosphaera watsonii* WH 8501 (17) (both unicellular night-time N_2_-fixers) and of *Trichodesmium erythraeum* IMS101 (a filamentous non-heterocystous day-time N_2_-fixer). We also compared expression to whole genome expression of *Prochlorococcus* sp. MED4 (18) (a marine non-N_2_-fixer) in order to determine how UCYN-A gene expression compares to general cyanobacterial gene expression in a sympatric open ocean species. We found that many genes in UCYN-A have distinct diel expression patterns and that UCYN-A has unusual gene expression patterns in comparison to unicellular N_2_-fixing cyanobacteria that fix N_2_ in the dark; however, it shares some general patterns with daytime N_2_-fixing cyanobacteria, with heterocysts of heterocyst-forming cyanobacteria and with unicellular non-N_2_-fixing cyanobacteria. Results suggest that optimal metabolism for open ocean cyanobacteria is aligned to the light period, and that symbiosis has enabled the unicellular UCYN-A to shift N_2_ fixation to the daylight period.

## Results and discussion

### UCYN-A has a daily rhythm of gene transcription

UCYN-A has clear diel patterns of gene transcription, with a large fraction of genes that had periodicity of transcript levels over the dark and light periods (27%).

About a third of the UCYN-A genome (31% genes) targeted by the array were transcribed at detectable levels (365 of 1194 total genes in UCYN-A1 and 394 of 1244 total genes in UCYN-A2, respectively) (Table S1). Approximately 85% of these genes have differences in transcript levels between dark and light periods, accounting for 27% of the total genes in each strain (Table S1 and S2). *C. watsonii, Cyanothece* sp. and *Trichodesmium* cultures also had a large fraction of genes with changes in the transcript levels between dark and light periods (39% in *C. watsonii*, 20% in *Cyanothece* sp. and 34% in *Trichodesmium*) (Tables S1 and S2).

The UCYN-A transcription values (log_2_-transformed) ranged from 2 to 13.5 with the median of 6.0. In both sub-lineages, genes coding for nitrogenase (*nif*), F_0_F_1_-ATP synthase (*atpA, atpB*), cytochrome *b_6_f* complex (*petB*, *petC, petF, petL*) and the photosynthetic gene *psaC* were the most highly transcribed in comparison to all detected genes (Table S5). Transcript levels of the same genes were also high for both sub-lineages in metatranscriptomes collected during the TARA expedition in the South Atlantic Ocean (19).

The two UCYN-A sub-lineages had similar periodicity of transcript levels to each other, despite divergence in gene sequences at the amino-acid level (average 14% genome-wide), cell morphology (19) and genome size (Figure 1). There were four gene clusters based on the time of day exhibiting the highest relative transcript level (Figure 1). Cluster I had the highest relative transcript level during the day (with a maximum 10 h into the light period) and included genes involved in cell division (e.g. *ftsZ, murG, minE, murB*), DNA replication (e.g. *topA, rpoE, DPO3B*), ABC transporters (e.g. *nikA, nikB, pstC, cbiO*), carbohydrate and lipid metabolism (e.g. *pdhA, pgi, fabG, fabH*) and a few photosynthesis genes (*petL, psaD and ccsB*). The transcripts for the *petL* gene, encoding subunit 6 of the cytochrome *b_6_f* complex and the only nitrogen fixation-related gene in this cluster (*nifK*) had a substantial change at this time (more than 3-fold).

**Figure 1.**
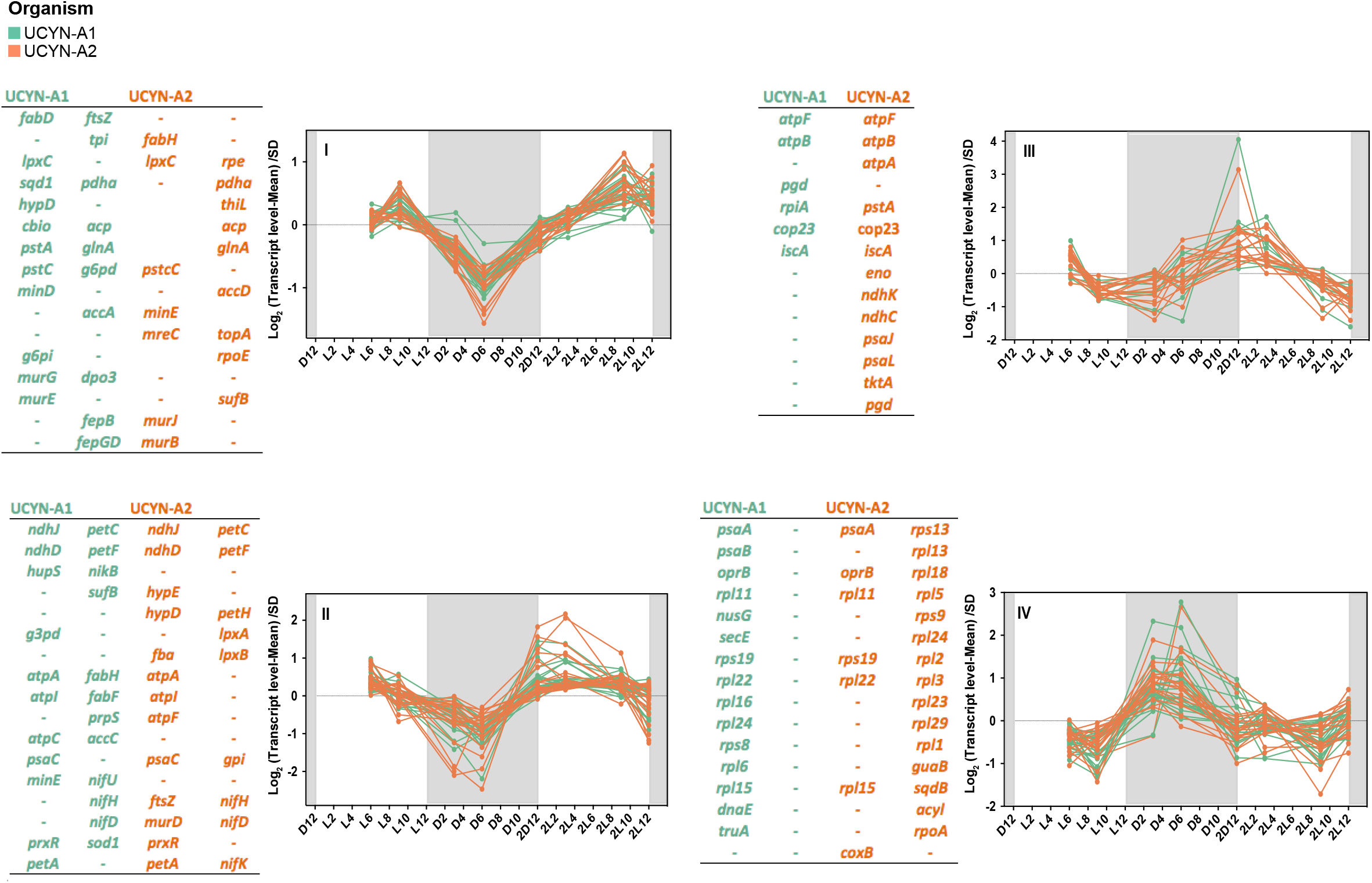
Four different clades based on Pearson correlation of the transcription profile of UCYN-A1 and UCYN-A2 genes over light-dark cycles. The transcription value of each gene at each time point was normalized to the mean at all time points and divided by standard deviation (SD) (*Y* axis, log 2 scale). The *X* axis represents time points where D and L stand for dark and light, respectively, followed by the corresponding hour into the light or dark periods. The second light-dark cycle is shown as 2D followed by the number of the corresponding hours entering light or dark period. The shaded area represents the dark period. In each cluster, most representative genes are listed in the table attached to the plot. UCYN-A1 genes are coded in green and UCYN-A2 genes are coded in orange.

The transcript abundance of genes from clusters II and III had similar patterns, with an increase before sunrise and a decrease during the dark period. The highest relative transcript levels for clusters II and III were 4h and 1h after sunrise, respectively, and included genes involved in nitrogen fixation (*nifHDK* operon) that increased 4-fold during the light period. However, these clusters also included genes involved in oxidative phosphorylation (e. g., NADH dehydrogenases subunits and ATP synthase related genes), carbohydrate catabolism such as those involved in glycolysis (e.g. *gap1, fbaA, pgi, eno*), the pentose phosphate pathway (*opcA* and *zwf*) and photosynthesis (e.g. cytochrome *b_6_f* complex subunit genes). In most cyanobacteria, genes encoding proteins involved in carbohydrate catabolism are highly transcribed during the night and are essential for survival under dark conditions.

The gene with the most dramatic difference in transcript levels between the light and dark periods encoded the membrane protein COP23 (23 kDa circadian oscillating protein), which had more than a 5-fold change in transcript abundance in both UCYN-A strains (Figure 1). COP23, a protein which may have a critical role in membrane function, has only been detected in nitrogen-fixing cyanobacteria (20).

Cluster IV had genes with the highest transcript level during the night and lowest during the day and included genes encoding photosystem I (PSI) subunits, a carbohydrate porin (*oprB*) and also genes encoding ribosomal proteins with 2- and 4-fold changes during the night period. Cluster IV had the lowest number of genes compared with the other clusters. Surprisingly, the PSI genes (*psaA* and *psaB*) were expressed during the night as in many anoxygenic phototrophic bacteria (21), whereas in most oxygenic cyanobacteria (including mats) these genes are expressed during the day (22).

The results show that UCYN-A has a daily rhythm of gene expression with strong periodicities of transcript levels over the diel cycle. Daily patterns of gene transcription in cyanobacteria are typically regulated by a circadian rhythm mediated by *kai* gene products (11). Rhythmic daily transcription patterns are still possible without the full suite of *kai* genes, for example, the marine cyanobacterium *Prochlorococcus* sp. MED4 lacks one of the circadian genes, *kaiA*, yet it maintains strong diel gene transcription patterns (18). However, *Prochlorococcus* sp. PCC 9511 loses the typical periodicities of the circadian clock under continuous light (23). In the case of UCYN-A, it lacks two of the three *kai* genes (24), which is unique among cyanobacteria, and furthermore, the *kaiC* gene was not transcribed at detectable levels. It is unclear what controls the UCYN-A diel gene expression pattern, but it could be that 1) there are unidentified components of a clock and signal transduction pathway, or that 2) the pattern could be driven by the physiological differences between light and dark conditions, which might be primarily driven by energy supplied by the eukaryotic partner. It is possible that the diel transcription patterns in UCYN-A are primarily regulated by the daily host metabolism, which itself is likely to be circadian. However, it is not yet known whether the UCYN-A diel cycle is maintained under constant conditions in UCYN-A, or whether the diel pattern is maintained in the absence of the partner alga.

### UCYN-A transcription patterns are similar to aerobic marine daytime N_2_-fixers and non-N_2_-fixers

UCYN-A had diel whole genome expression patterns that were different from those of phylogenetically closely related unicellular cyanobacteria (17). Only a few genes (such as those encoding ATP synthase) had the same daily pattern among all cyanobacteria, presumably differing because of physiology (e.g. N_2_-fixing or not). The unicellular cyanobacteria *C. watsonii* WH 8501 and *Cyanothece* sp. ATCC 51142, which fix N_2_ during the night, expressed many genes in an opposite pattern compared to the day-time N_2_-fixing *T. erythraeum* and UCYN-A (Figure 2 and Tables S4 and S5). Interestingly, the diel transcription patterns of N_2_ fixation and PSI genes in UCYN-A were opposite to those in *Cyanothece* sp. ATCC 51142 and *C. watsonii* WH 8501 and more similar to those of *T. erythraeum* (Figure 2 and Tables S4 and S5).

**Figure 2.**
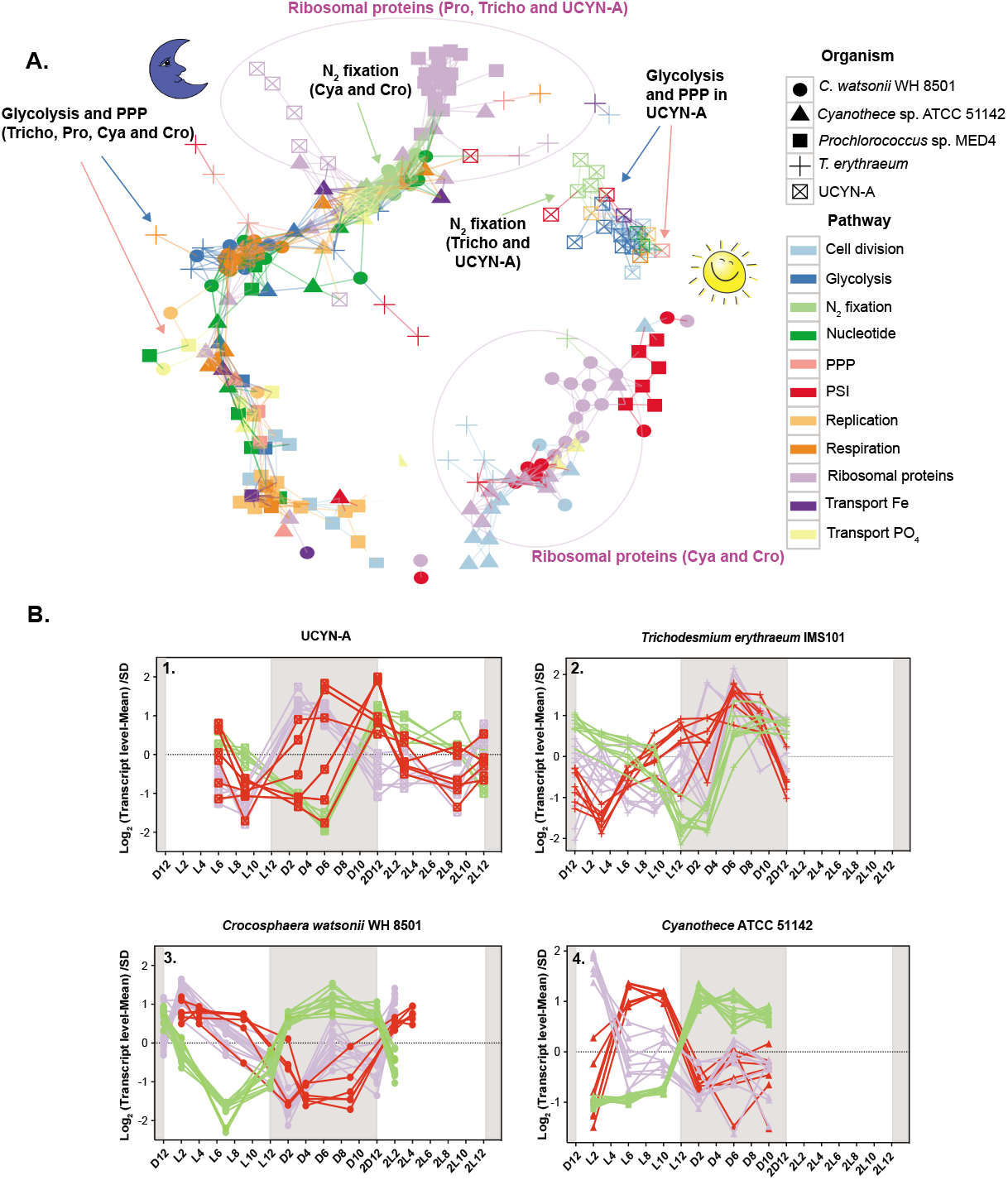
**A.** Transcriptional network based on Pearson correlation of gene transcription over the diel cycle in all studied cyanobacteria. The genes are connected if correlation coefficient for their transcription patterns is higher than 0.5. The genes shown are diel genes with variable transcription patterns among the studied cyanobacteria. The arrows point to genes for glycolysis, PPP and N_2_ fixation in the studied diazotrophs. The purple circles demarcate genes for ribosomal proteins included in the analysis. Abbreviations: *Prochlorococcus* sp. MED4 (Pro), *Cyanothece* sp. ATCC 51142 (Cya), *C. watsonii* WH 8501 (Cro), *T. erythraeum* (Tricho), Pentose Phosphate Pathway (PPP), Photosystem I (PSI). **B.** Four time course plots are attached for the N_2_-fixing cyanobacteria showing the diel transcription patterns of photosystem I, N_2_ fixation and genes for ribosomal proteins.

As observed for the activity of nitrogenase, it has been demonstrated that levels of *nif* transcripts and the biosynthesis of different components of the nitrogenase complex are very sensitive to O_2_ (22, 25–27), most likely to avoid energy losses associated with the degradation of this enzyme by O_2_. Thus, the different patterns observed in the genes involved in N_2_-fixation in the cyanobacteria studied here presumably are due to the different mechanisms used to protect the nitrogenase complex from the O2 produced by photosynthesis. *T. erythraeum* and UCYN-A had the maximum transcript levels of the nitrogenase and PSI genes just prior to dawn, but maintained high levels of transcripts for both sets of genes during the day. The peak of transcript levels just before dawn is likely due to the advantage of synthesizing nitrogenase in preparation for N_2_ fixation in the early hours of the day (28).

The diel expression patterns of genes that are unrelated to N_2_ fixation in the aerobic day-time N_2_-fixers (*T. erythraeum* and UCYN-A) were also more similar to those of non-N_2_-fixing sympatric cyanobacteria of the genus *Prochlorococcus* and to heterocysts of heterocyst-forming cyanobacteria than to the nighttime N_2_-fixing cyanobacteria (C. *watsonii* and *Cyanothece* sp.). The transcript levels of genes encoding ribosomal proteins in both UCYN-A and *T. erythraeum* were higher during the night, probably because the reduced nitrogen required for the synthesis of new proteins was obtained during the day (Figure 2 and Tables S4 and S5). Similar patterns were observed in *Prochlorococcus* with higher transcript levels during the night (Figure 2 and Tables S4 and S5) while genes encoding ribosomal proteins in *C. watsonii* WH 8501 and *Cyanothece* sp. ATCC 41142 had maximum transcript levels during the day (Figure 2 and Tables S4 and S5). Intriguingly, these results imply that both UCYN-A and *T. erythraeum* have adopted day-time gene transcription patterns for the main metabolic pathways minimizing cellular processes in the dark. The night-time patterns of the transcript levels of the ribosomal proteins (genes) would make it possible to have proteins synthesized in order to make the most efficient use of the light period, as in *Prochlorococcus*. Because UCYN-A and *Trichodesmium* are likely to be the two most abundant N_2_-fixing cyanobacteria in the open ocean, it appears that direct coupling of N_2_ fixation to photosynthesis is important in the oligotrophic environment (as long as low oxygen concentrations are maintained in the cell).

Phosphorus is a vital element for cellular energetics and growth and is acquired by oceanic bacterioplankton primarily as phosphate (29–31). The UCYN-A phosphate ABC transporter had the same diel pattern as in *Trichodesmium* for genes involved in DNA replication, with higher transcript levels during the day (Table S5), but maximum transcript abundances during the late afternoon in *Crocosphaera* and *Cyanothece* (17, 32). High levels of phosphate transporters during the day could meet the increased demand for inorganic phosphate (33, 34) during DNA replication, which occurs during the day in UCYN-A and *Trichosdesmium*. Similar patterns were observed in the heterocyst-forming *Richelia* with peak expression of P acquisition genes at approximately 15:00, suggesting the apparent rhythmicity of P acquisition could be a common feature of daytime N_2_-fixers (35).

The initiation factor of DNA replication, DnaA, is a protein highly conserved in prokaryotes although it is absent in red algae, the cyanobacterial symbiont *Nostoc azollae* (36) and also the spheroid bodies of diatoms (37). The genome of UCYN-A lacks the *dnaA* gene as well. Recent studies suggested that DnaA is not essential for DNA replication and the lack of *dnaA* could suggest a preadaptation of the genome to enable the symbiosis (38). In UCYN-A and *T. erythraeum*, genes for DNA replication (*dnaE and* RNaseHI), DNA topoisomerases, DNA gyrases and cell division (*ftsZ*, *mre, min*) had maximum transcript levels during the day (i.e., after midday), and minimum levels at night (Figure 3A and Figure S1). In contrast, the nighttime N_2_-fixing *Cyanothece* sp. ATCC 51142 and *C. watsonii* WH 8501 confine cell division to the period of transition from dark to light at sunrise. The temporal delay in cell division in *Cyanothece* and *Crocosphaera* has been suggested to reflect the need to recover energy reserves with light-derived energy after night-time metabolic activity (39). The similarity of the pattern in UCYN-A to *Trichodesmium* is consistent with UCYN-A shifting metabolism to the daytime.

**Figure 3.**
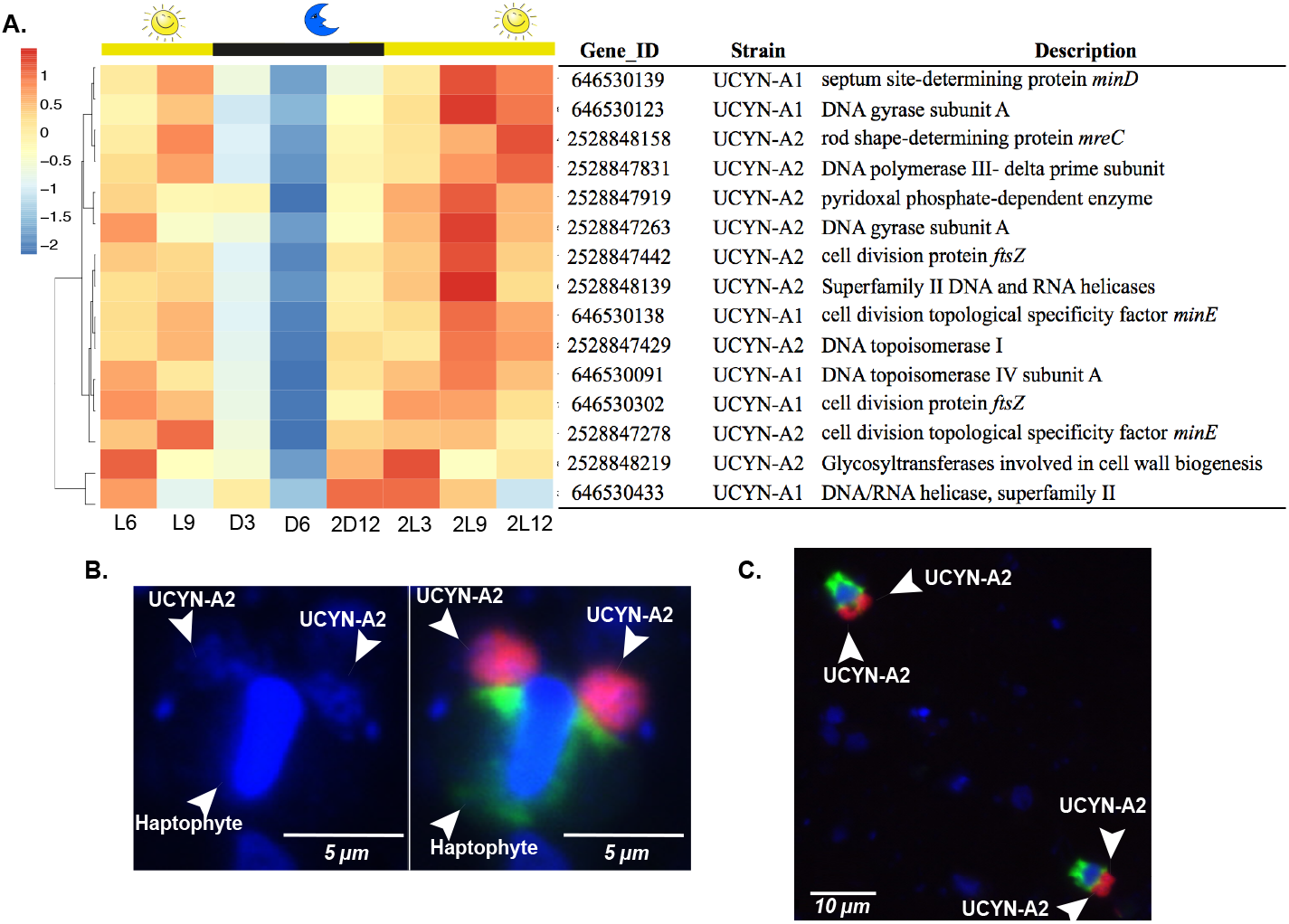
Transcription of genes for replication and cell division in UCYN-A. **Upper Panel: A)** Diel transcription patterns for cell division and replication genes in UCYN-A1 and UCYN-A2 over the light-dark cycle. Hierarchical clustering of genes was based on Pearson correlation between their transcription profiles. The transcription values of genes at each time point were standardized, and the blue-red scale shows how many standard deviations a transcription value was lower or higher, respectively, from the mean transcription values over the diel cycle (Z score). Gene ID and gene product corresponding to each gene for UCYN-A1 and UCYN-A2 are shown. Time is shown on X-axis as light (L) and dark (D), respectively, followed by the corresponding hour after the sunrise and sunset periods started. The second light-dark cycle is shown as 2D followed by the number of the corresponding hours entering light or dark period. **Lower Panel:** Epifluorescence micrographs of dividing UCYN-A2 detected with CARD-FISH (19). **B)** Two big clusters of UCYN-A2 cells and the haptophyte host attached. Left Panel: the nucleus of the host and the UCYN-A2 cells were visualized with–DAPI stain (blue). Right Panel: The UCYN-A2 (red) and its haptophyte host (green). **C)** Two different associations of UCYN-A2 with its haptophyte dividing in samples from Scripps Pier.

Microscopy counts of the *B. bigelowii* -UCYN-A2 symbiosis were performed eight times during two diel cycles in order to observe the timing of cell division (Figures 3B and C and Table S6). In both diel cycles, single host cells with two associated UCYN-A2 cells (or groups of cells), corresponding to approximately 60% of total cell counts, were present at night between 21:00 and 03:00. The delay observed between the higher transcription levels after midday and actual cell division at 21:00 may be explained by the need of the cell to coordinate the assembly of the cell division machinery prior to cell division.

### Unique UCYN-A transcription patterns

Although many gene transcription patterns in UCYN-A are more similar to *Trichodesmium* than to other unicellular N_2_-fixing cyanobacteria, some of the patterns were unique to UCYN-A. Such unique gene transcription patterns in the UCYN-A symbiosis may provide clues to possible roles of specific genes involved in adaptation to N_2_-fixing symbiosis revealing metabolic interdependence between host and symbiont. In order to compare the transcriptomic patterns of these specific genes with the rest of the N_2_-fixers, we performed network analysis of these genes using Pearson correlation. Whereas most of the key genes of the major pathways in UCYN-A had higher transcript levels during the day, the other unicellular N_2_-fixing cyanobacteria had maximum transcript levels at night (Figure 4). For example, glycolysis genes in UCYN-A had the highest levels of transcripts at sunrise and midday (maximum light conditions) in contrast to the other cyanobacteria (Figures 4 and 5). The metabolic pathway that generates reductant for biosynthesis activities (NADPH), the pentose phosphate pathway (PPP), had similar patterns. The allosteric effector *opcA*, which redirects carbon flow to the first enzyme of the PPP (glucose-6-P dehydrogenase (*zwf)*) (18, 40), had a different periodic transcript level pattern in UCYN-A (Figures 4 and 6) compared to other cyanobacteria (41, 42).

**Figure 4.**
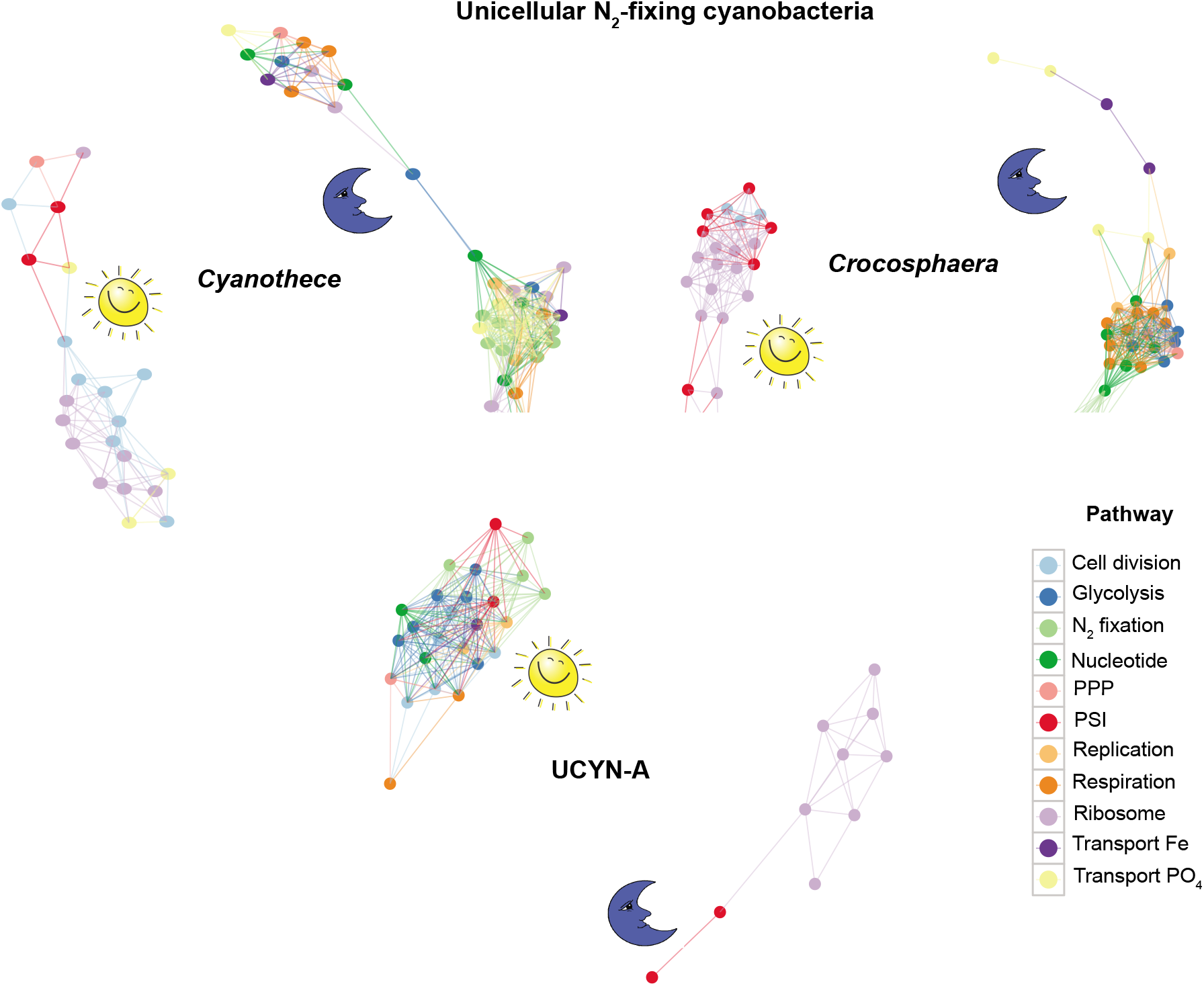
Network showing the Pearson correlation for gene transcriptions in the unicellular N_2_-fixing cyanobacteria *Cyanothece* sp. ATCC 51142 (*Cyanothece), C. watsonii* WH 8501 (*Crocosphaera*) and UCYN-A. Shown here are key genes in major metabolic pathways with distinct diel transcription patterns. The genes are connected if their correlation coefficient for transcription patterns is higher than 0.2. PPP, pentose phosphate pathway; PSI, photosystem I.

**Figure 5.**
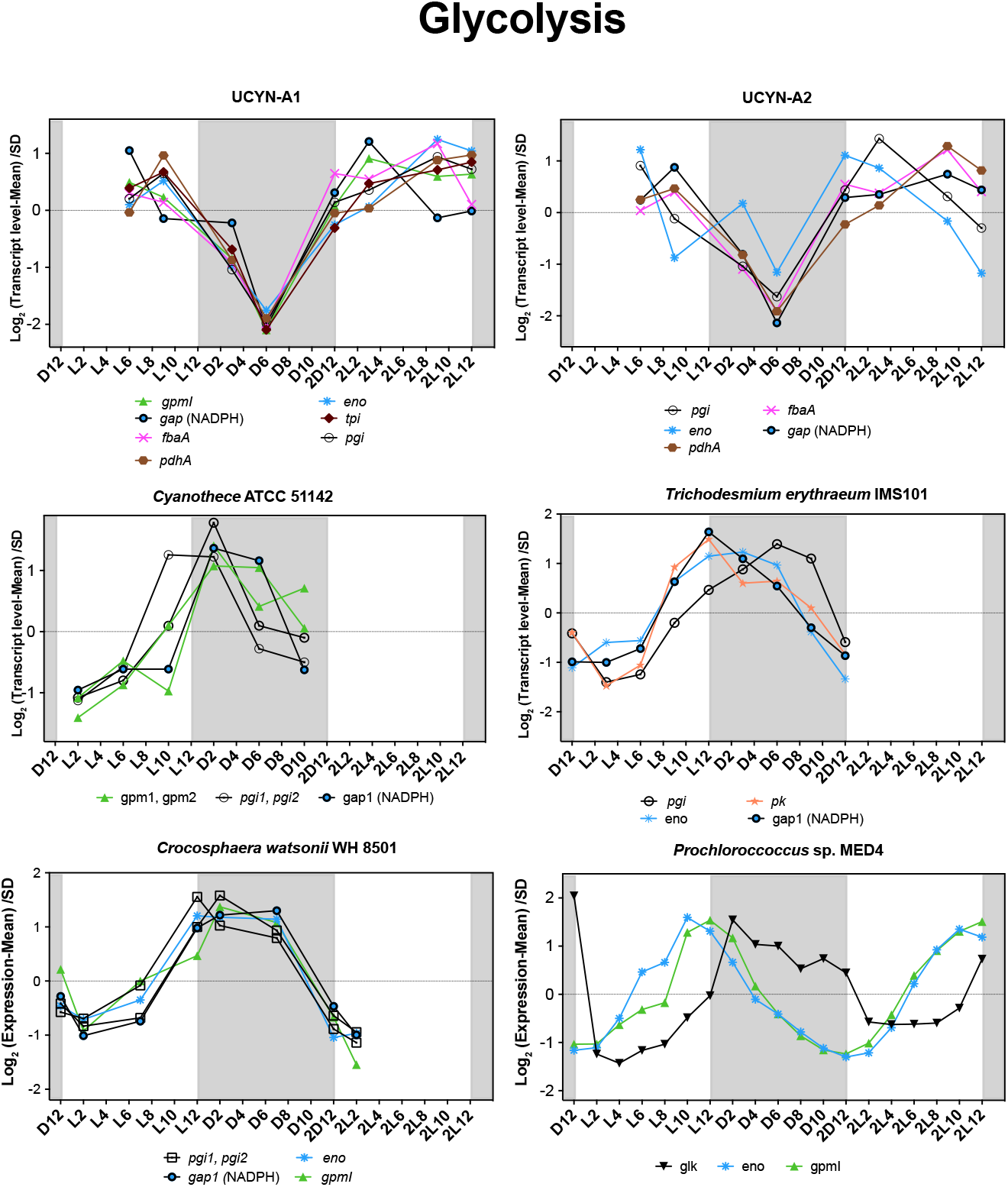
Transcriptional profiles of the genes for glycolysis over light–dark cycles in the cyanobacteria studied here. The transcription value of each gene at each time point was normalized to the mean at all time points and divided by standard deviation (SD) (*Y* axis, log scale). The *X* axis represents time points where D and L stand for dark and light, respectively, followed by the corresponding hour into the light or dark periods. The second light-dark cycle is shown as 2D followed by the number of the corresponding hours entering light or dark period. The shaded area represents the dark period.

**Figure 6.**
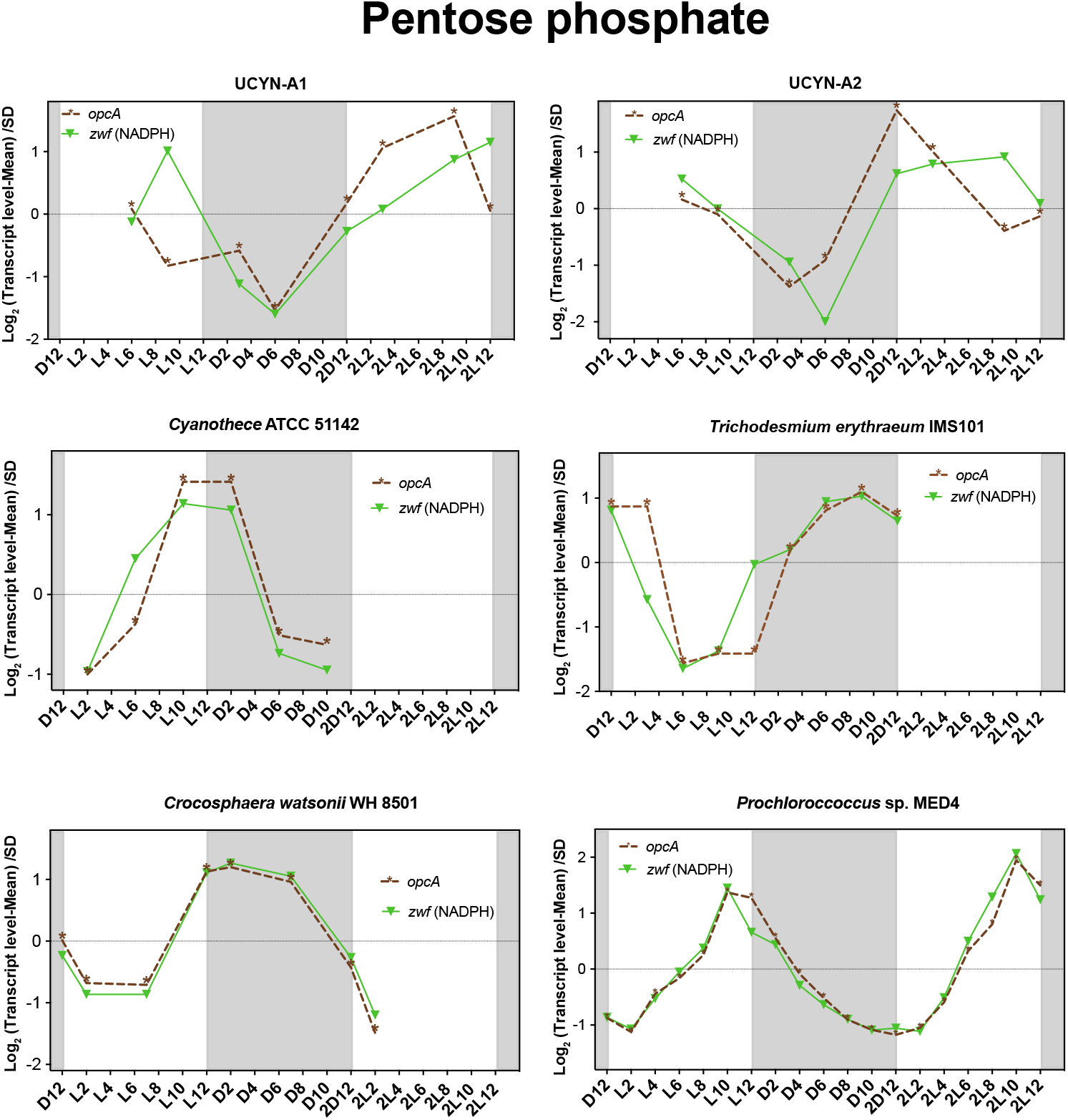
Transcriptional profiles of *opcA* (allosteric effector) and *zwf* (glucose-6-P dehydrogenase) over light–dark cycles in the cyanobacteria studied here. The transcription value of each gene at each time point was normalized to the mean at all time points and divided by standard deviation (SD) (Y axis, log scale). The *X* axis represents time points where D and L stand for dark and light, respectively, followed by the corresponding hour into the light or dark periods. The second light-dark cycle is shown as 2D followed by the number of the corresponding hours entering light or dark period. The shaded area represents the dark period.

N_2_ fixation in UCYN-A depends on the light period for the supply of photosynthate from the host during the day, as well as possibly producing ATP by cyclic photophosphorylation with PSI. Because UCYN-A cannot fix carbon dioxide, it has to obtain reduced carbon compounds in the same way. Based on genome and transcriptomic profiles, we propose a pathway of carbon metabolism for the regeneration of reductant and ATP in UCYN-A, which is needed for N_2_ fixation (Figure 7). Carbohydrate porins or ABC transporters could transport the carbohydrates from the host to the cyanobacteria during the day and the carbon compounds metabolized through the oxidative pentose phosphate (OPP) or glycolysis pathways. Pyruvate is required for generation of reductant for nitrogenase and also to generate acetyl-CoA for synthesis of fatty acids.

**Figure 7.**
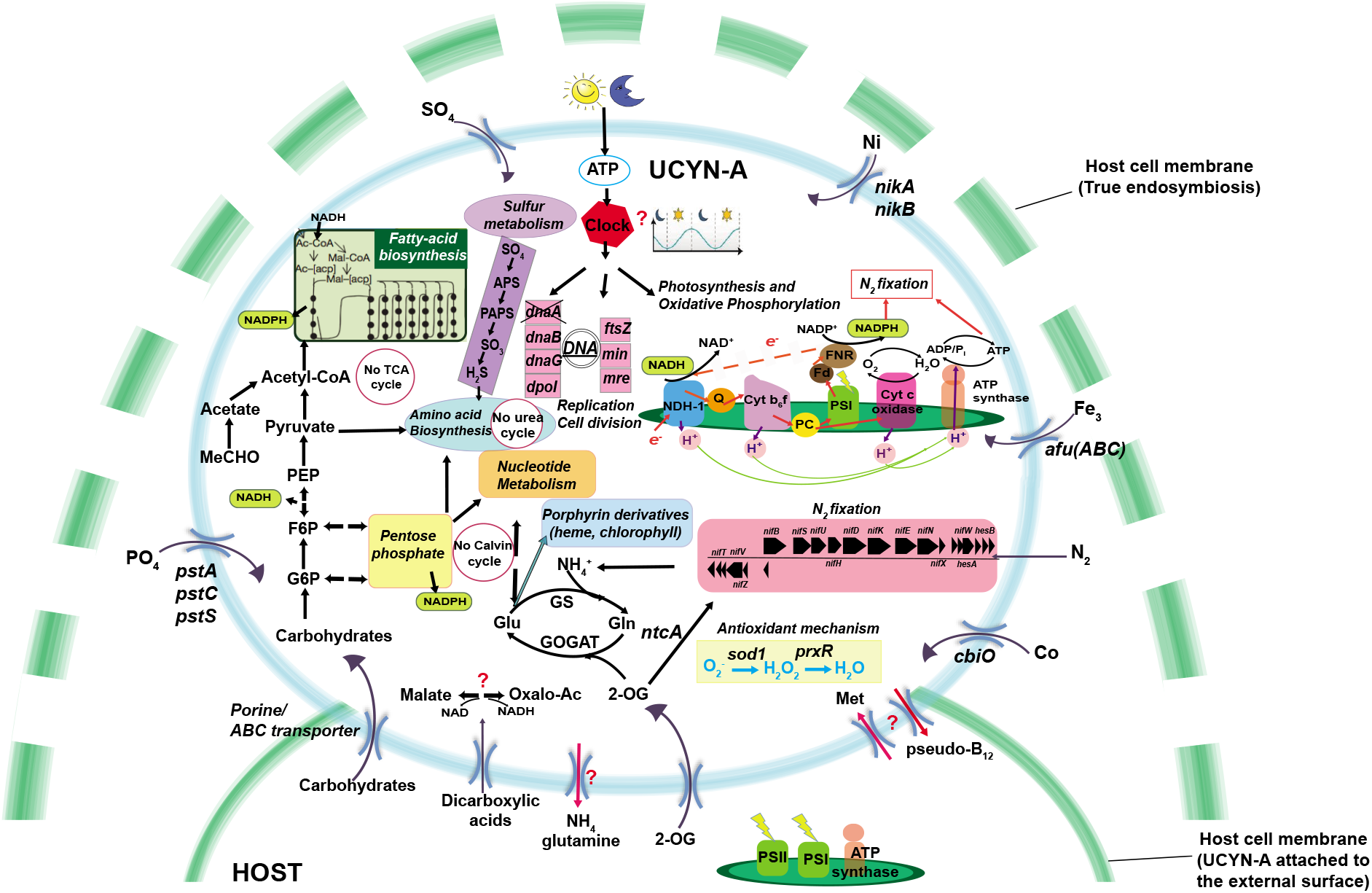
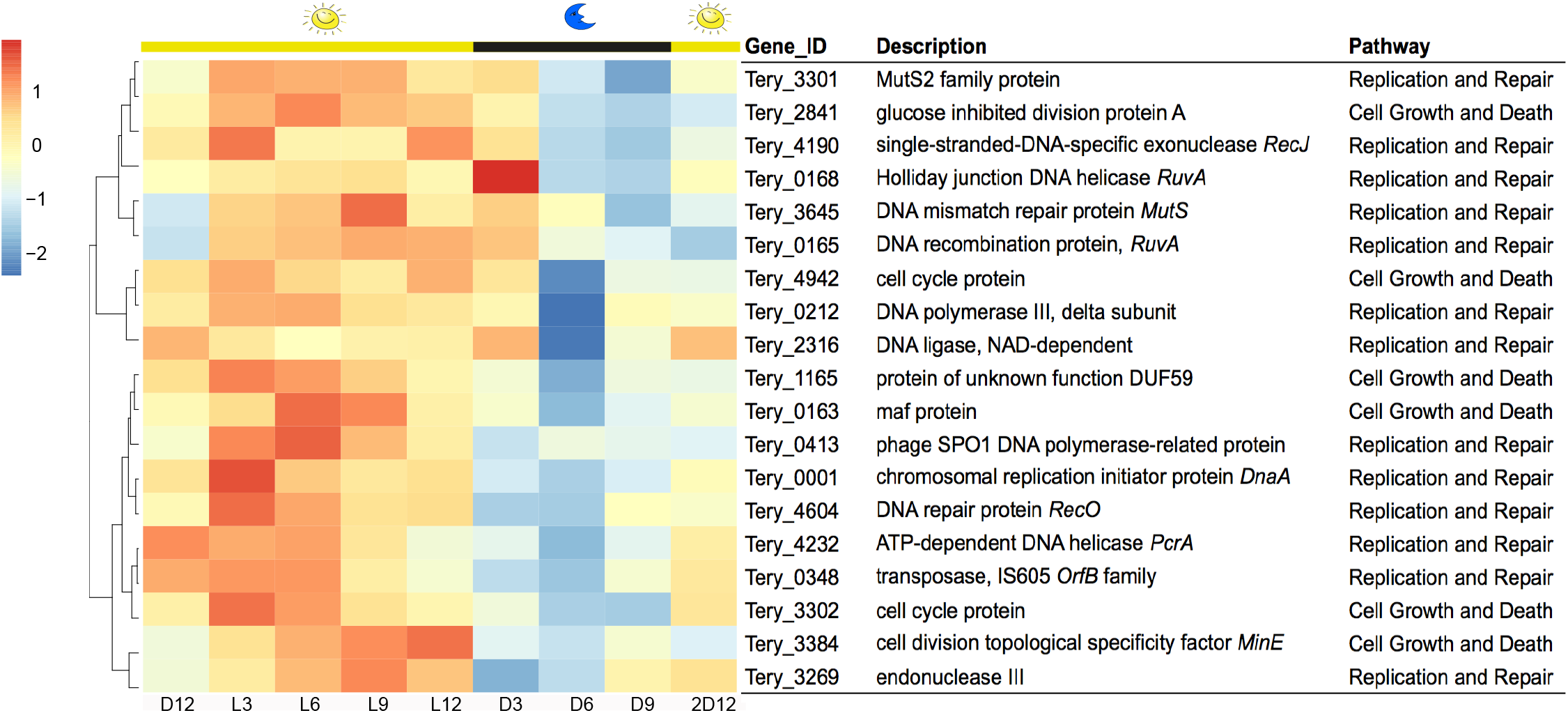
Schematic model of UCYN-A showing the possible main cellular functions, metabolic pathways and transporters.

Because UCYN-A lacks photosystem II, which normally supplies electrons to photosystem I by splitting water, UCYN-A needs alternative electron donors if it uses PSI to make the reductant NADPH. The NADH generated by the OPP pathway or by glycolysis could reduce the plastoquinone pool via the NDH-1 complex and transfer electrons to ferredoxin though the PQ pool, cytochrome *b_6_f* plastocyanin and the action of PSI. Ferredoxin could deliver electrons to the ferredoxin:NADPH oxidoreductase (FNR), which might provide reductant and ATP directly to the dinitrogenase reductase. To increase the ATP/e^-^ ratio, UCYN-A can redirect electrons from PSI to NDH-1 in cyclic phosphorylation. This mechanism to supply nitrogenase with electrons was proposed years ago for heterocysts (43).

Together, the results are consistent with the assumption that UCYN-A uses host-supplied carbohydrates during the day while other unicellular cyanobacteria synthesize their own carbohydrates during the day and use them during the evening or at night. The unique distribution of these metabolic processes suggests that UCYN-A has developed the ability for light-driven, day-time N_2_ fixation under oxic conditions as a result of symbiosis.

Apart from fixed carbon, several other compounds may be made available to UCYN-A, which may be endosymbiotic, and relies on the host for all of its essential nutrients. Interestingly, UCYN-A has the whole pathway for the synthesis of the cyanobacterial type of vitamin B_12_, pseudocobalamin, that can be required for the activity of several vital enzymes in central metabolism (44) (Table S8). Transcription of genes involved in B_12_ synthesis were detected in all cyanobacteria, and some of them had diel patterns (Table S2 and S8). It is unknown if UCYN-A has enzymes that require pseudocobalamin or whether it can be used by the host. However, in order for the host to use pseudocobalamin, it would have to be remodeled in order to be accessible to the haptophyte (45). The role of pseudo-B_12_ biosynthesis in UCYN-A is unclear, but the fact that UCYN-A retains this entire pathway, in such a reduced genome, indicates that it is likely to have an important role, perhaps in symbiosis.

It is still unclear how N_2_ fixation in UCYN-A avoids the oxygen evolved by the photosynthetic host alga. There are only two possible pathways for consuming O2 in UCYN-A, including aerobic (cytochrome-dependent) respiration and the photocatalyzed reduction of O2 to H_2_O in PSI which occurs in the heterocysts of cyanobacteria like *Nostoc* sp. PCC 7120 (46–48). The latter, called the Mehler reaction, results in the production of the superoxide radical O_2_^-^, which is subsequently reduced to water (49, 50).

In UCYN-A, the cytochrome *c* oxidase *coxA* gene was transcribed during the night (cluster IV) but also rarely during the day, along with a few N_2_ fixation genes (cluster I) (Figure 1). Moreover, we also found higher transcript levels during the day for the antioxidant enzyme superoxide dismutase (*sod1*) and two peroxiredoxins (*prxR*), which have the ability to detoxify peroxide (Figures 1 and 7). Both antioxidants would protect the nitrogenase against the reactive oxygen species produced by UCYN-A or the haptophyte host (Figure 7).

It is not currently possible to directly determine the oxygen protection mechanisms in this uncultured microorganism because 1) transcription cannot necessarily be related to function and 2) it is not possible to do physiological experiments with this low-abundance microorganism that has yet to be obtained in an axenic culture. Consequently, the question of protection from O_2_ cannot be directly addressed experimentally, but our results suggest that some of the proteins in UCYN-A could help to protect nitrogenase from the O2 generated by host photosynthesis.

Because UCYN-A has genome reduction normally associated with endosymbiosis (e.g. in *Paulinella chromatophora*; (51)), the unique gene transcription patterns of UCYN-A may provide insights into the evolution of endosymbiosis and organellar evolution. Future studies are needed to determine if the rhythm of these patterns is maintained under constant conditions as in a circadian rhythm, whether the host has a circadian rhythm and/or the daily cycle in UCYN-A simply responds to metabolite availability from the host. It will also be interesting to determine how PSI is involved in supporting the energy or reductant requirements of N_2_ fixation. Such experiments will have to await the establishment of a pure culture.

## Materials and Methods

### Diel sampling of UCYN-A

Surface seawater samples for UCYN-A transcription and catalyzed reported deposition-fluorescence *in situ* hybridization (CARD-FISH) analyses were collected using a bucket from the end of the Scripps Institution of Oceanography (SIO) Ellen Browning Scripps Memorial Pier in La Jolla, CA, USA. Two replicates were collected from the bucket at each time point within 48 hours between 28^th^ July and 1^st^ August 2014 for transcriptomic analysis and between 3^rd^ - 8^th^ May 2016 for CARD-FISH. A total of 16 samples were collected every 3-6 hours (two replicates taken at each of eight time points): 12:00-L6, 15:00-L9, 21:00-D3, 00:00-D6, 06:00-2D12, 09:00-2L3, 15:00-2L9 and 18:00-2L12. L and D stand for light and dark period, respectively, 2L and 2D the second light-dark cycle, and the number the corresponding hours entering light or dark period.

For the CARD-FISH assay, from each seawater replicate, 190 mL of seawater was fixed with 10 mL 37% formaldehyde (1.87% v/v final concentration) at 4°C in the dark for 1 hour. After fixation, 100 mL was filtered at a maximum vacuum pressure of 100 mm Hg onto a 0.6 μm pore-size, 25 mm diameter polycarbonate membrane filter (Millipore Isopore™, EMD Millipore, Billerica, MA, USA) with a support filter of 0.8 μm pore-size, 25 mm diameter polycarbonate cellulose acetate membrane filter (Sterlitech Corporation, Kent, WA, USA). The filters were kept at −80°C until processed.

Samples for RNA extraction were collected by filtering a total of 500 mL from each seawater replicate through 0.22 μm pore-size, 47 mm diameter Supor filters (Pall Corporation, Port Washington, NY, USA) using a peristaltic pump. Filters were placed in sterile 2 mL bead-beating tubes with sterile glass beads, flash-frozen in liquid nitrogen and stored at −80°C until extraction.

### Double CARD-FISH assay

The double CARD-FISH assay was carried out following the protocol designed by Cabello et al. 2016 and Cornejo-Castillo et al. 2016. All of the probes, competitors and helpers used in this work are compiled in Table S7. More details are described in Supplementary Information. Microscopic evaluation and counting was performed with the Carl Zeiss Axioplan-2 Imaging Fluorescent Microscope (Zeiss, Berlin, Germany) in 3 transects (8.0 x 0.1 mm^2^ each) across the filter piece. Cell dimensions were estimated using AxioVision 4.8 and Image J software (52).

### Diel sampling of Trichodesmium erythraeum IMS101 cultures

Biological triplicate cultures of *T. erythraeum* were grown in rectangular canted neck polycarbonate cell culture flasks with a 0.2 μm pore-size vent cap and 225 cm^2^ surface area (Corning Inc., Corning, NY, USA). The cultures were maintained at 26°C on a 12h:12h light:dark cycle at 50 μmol quanta m^-2^ s^-1^ in YBCII media (53) supplemented with 2.8 μmol L^-1^ ferric ammonium citrate. The light was set on at 7:00 and off at 19:00 hours. The cultures were 10-fold diluted from the inoculum and were verified to be axenic by staining with DAPI and visualizing cells under an epifluorescence microscope (Carl Zeiss, Thornwood, NY, USA). Growth and cell density were monitored until the cultures reached exponential phase (~10-14 days after inoculation), during which the cells were harvested for the diel transcription assay. Samples were taken at 3 hours intervals starting at the onset of the light period until the end of the dark period for a total of 24 hours. A total of 27 samples were collected from these nine time points: 7:00-D12, 10:00-L3, 13:00-L6, 16:00-L9, 19:00-L12, 22:00-D3, 1:00-D6, 4:00-D9 and 7:00-2D12, where L and D stand for light and dark period, respectively, 2D the second light-dark cycle, and the number the corresponding hours entering light or dark period. At each time point, 200 mL each of triplicate cultures (replicates from different flasks) was filtered onto a 5 μm pore-size, 47 mm diameter polycarbonate membrane filter (Osmonics, Minnetonka, MN, USA). The filters were immediately frozen in liquid nitrogen and stored at −80° C until processing.

### RNA extraction and processing for hybridization to the microarray

Environmental RNA containing transcripts from UCYN-A cells was extracted using the Ambion RiboPure Bacteria kit (Ambion®, ThermoFisher), with modifications that included mechanical lysis using glass beads (Biospec, Bartlesville, OK). The extracted RNA was treated with Turbo-DNA-free™ DNase Kit (Ambion®, ThermoFisher) to remove genomic DNA. Sufficient environmental RNA was obtained for two replicates at 4 sampling times (L6, L9, D3 and 2L12): L6-1, L6-2, L9-1, L9-2, D3-1, D3-2, 2L12-1 and 2L12-2. L and D stand for light and dark period, respectively, 2L and 2D the second light-dark cycle, and the number the corresponding hours entering light or dark period.

Total RNA for *T. erythraeum* was extracted using the Ambion RiboPure Bacteria kit (Ambion®, ThermoFisher), followed by in solution DNase digestion with the RNase-free DNase kit and on-column cleanup with the RNeasy MiniElute kit (Qiagen, Valencia, CA, USA).

RNA purity, concentration and quality were determined using a NanoDrop 1000 (Thermo Scientific, Waltham, MA, USA) and a 2100 Bioanalyzer (Agilent Technologies, Santa Clara, CA, USA) using the RNA 6000 Nano kit (Agilent Technologies). Only samples with RNA Integrity Number >7.0 and ratios of A260/A230 and A260/A280 ≥1.8 were processed further.

From environmental RNA samples that contained UCYN-A, double-stranded (ds) cDNA was synthesized and amplified following the procedure described in Shilova et al..(54). Briefly, 400 ng RNA from each sample was used, and 1 μL of 1:100 dilution (corresponding to 4.7 attomoles of ERCC-0016) of the (External RNA Control Consortium, (55)) RNA spike-in mix 1 (Ambion®) was added before amplification to monitor the technical performance of the assay (55).

Double-stranded cDNA was synthesized and amplified using the TransPlex Whole Transcriptome Amplification kit (WTA-2, Sigma-Aldrich, St Louis, MO, USA) and antibody-inactivated hot-start Taq DNA Polymerase (Sigma-Aldrich). The amplified cDNA was purified with the GenElute PCR cleanup kit (Sigma-Aldrich), and the quality and quantity of ds-cDNA was determined with NanoDrop 1000 and a 2100 Bioanalyzer using the Agilent DNA 7500 kit (Agilent Technologies). Four hundred ng of total RNA yielded on average 12 μg of ds-cDNA. The labeling and hybridization of cDNA samples (1.0 μg of ds-cDNA) to the microarray was done at Roy J. Carver Center for Genomics (CCG) Facility (University of Iowa, Iowa city, Iowa, USA) according to the Agilent Technology for arrays protocol.

For *T. erythraeum*, at least 30 μg of unamplified total RNA with a concentration of 1.0 μg μL^-1^ per sample was provided for 27 samples. A control sample was generated by mixing equal amount of total RNA, based on NanoDrop measured concentration, from each of the 27 samples resulting in 28 samples in total. Reverse transcription of the total RNA, labeling of cDNA, and hybridization to the array were performed at the Roche NimbleGen facility according to the manufacturer’s protocol (Roche NimbleGen, Inc., Madison, WI, USA).

### Design of the UCYN-A array

The oligonucleotide expression array of UCYN-A was designed using UCYN-A1 and UCYN-A2 genes using eArray web-based tool (Agilent Technology Inc.; https://earray.chem.agilent.com/earray/) similar to the array design described in Shilova et al.(54). The gene sequences were obtained from the National Center of Biotechnology Information (NCBI, http://www.ncbi.nlm.nih.gov). Briefly, six probes of 60 nucleotides (nt) length were designed for each gene, and a total of 6618 probes (1199 genes) and 6862 probes (1246 genes) were designed for UCYN-A1 and UCYN-A2, respectively. These probes were replicated (4 times in the 8×60K array slides and 13 times in the 4×180K array slide) which allowed internal evaluation of signals. The sequences of all oligonucleotide probes were tested *in silico* for possible cross-hybridization as described below. The probe sequences were used as queries in the BLASTN against the available nt databases in June 2014: Marine microbes, Microbial Eukaryote Transcription and Non-redundant Nucleotides in the Community Cyberinfrastructure for Advanced Microbial Ecology Research and Analysis (CAMERA, http://camera.calit2.net/,(56)). Agilent technology allows 5% nt mismatch in the whole probe region, thus sequences with a range of 95-100% nt identity to the target probe are detected. Therefore, all probes with BLASTN hits with ≥95% over 100% nt length were deleted. Next, probe sequences that passed the cross-hybridization filter, were clustered using CD-HIT-EST(57, 58) at 95% nt similarity to select unique probes for UCYN-A1 and unique probes for UCYN-A2. Finally, to select probes specific for each strain, the probes with >95% nt identity to the genes in the other strain were deleted. However, a few probes that showed cross-hybridization between both strains for highly conserved genes (such as the nitrogenase gene, *nifH*) were retained. In summary, 6120 probes for 1194 genes of UCYN-A1 and 6324 probes for 1244 genes of UCYN-A2 were chosen.

In addition, standard control probes as part of the Agilent Technology Array (IS-62976-8-V2_60Kby8_GX_EQC_201000210 with ERCC control probes added) were included randomly to feature locations on the microarray slide. The final design of the microarray was synthesized on two platforms: ca. 62976 experimental and 1319 control probes on the 8×60K array slide and ca. 180880 experimental and 4854 control probes on the 4×180K array slide. The probe sequences are available at NCBI Gene Expression Omnibus (GEO) under accession number GSE100124.

### Design of the T. erythraeum IMS101 array

A custom oligonucleotide array for *T. erythraeum* was designed using the Roche NimbleGen platform: (NimbleGen design ID: 080610_Trich_erth_UCSC_TS_expr) according to the complete genome assembly of *T. erythraeum* IMS101 (NC_008312). The genome sequence is publically available via gateways including GenBank (https://www.ncbi.nlm.nih.gov/nuccore/NC_0083120), IMG (http://img.jgi.doe.gov:80/cgibin/pub/main.cgi?section=TaxonDetail&page=taxonDetail&taxon_oi_d=637000329), and UCSC genome browser (http://microbes.ucsc.edu/cgi-bin/hgGateway?db=tricEryt_IMS101). Up to six 60-nt long tiling probes were designed to target each of the 4788 genes in the genome, resulting in a total of 28235 probes. The probes were duplicated on the array to allow internal evaluation of hybridization signals. Moreover, tiling 60 nt oligonucleotide probes were also designed to target the intergenic regions >60 bp in length at a 150 bp interval, leading to a total of 11175 probes targeting 3877 intergenic regions (average 2.9 probes per intergenic region), however hybridization data for intergenic probes are not presented here. All the probes were rank ordered and selected based on the following criteria: 1) they must have a minimum annealing temperature of 68°C; 2) there is no cross contamination among the probes for different genes and for different intergenic regions. In addition to the experimental probes, standard control probes were also included on the microarray for quality assessment of the sample preparation, the hybridization process and the intensity measurements. The final microarray slides were printed in 4-plex (4×72K) format with 67645 experimental probe features and 7454 control probe features on one array. The full microarray platform descriptions and data for *T. erythraeum* are available at NCBI GEO under accession number GSE99896. Microarray hybridization signals were quantified using a GenePix 4000B Scanner (Molecular Devices, Sunnyvale, CA, USA) at the Roche NimbleGen facility.

### Microarray data analysis

All data analyses were performed with R (www.R-project.org) and the Bioconductor Project(59), specifically using the Biobase(60), Linear Models for Microarray LIMMA (61), arrayQualityMetrics(62), affyPLM(63, 64), and genefilter packages.

### 1) UCYN-A microarray

Transcription values for each gene were obtained using median polish summarization, and values were normalized using quantile normalization (63, 64) (Figure S2). The transcription values for UCYN-A at L6, L9, D3 and 2L12 are the mean transcription of the two replicates (L6-1, L6-2, L9-1, L9-2, D3-1, D3-2, 2L12-1 and 2L12-2). Raw and normalized microarray data for UCYN-A were submitted to NCBI GEO under accession number GSE100124. To determine if transcription of a gene was detected, the signal-to-noise ratio (SNR) of each chip was calculated as: SNR = (S_i_–BG)/BG; where S_i_ is the hybridization signal for the gene and BG is the chip background signal determined as average of the lowest 5% of all signals. Transcription was considered detected if SNR of a transcript was ≥5 (as in (Shilova et al. 2014) Transcription values were centered and scaled across genes and samples, and a distance matrix was calculated using Pearson’s correlation coefficient. The distance matrix was then used in hierarchical clustering by a complete agglomeration method to identify clusters of genes with similar patterns of transcription during the diel transcription.

### 2) T. erythraeum microarray

The raw microarray data for *T. erythraeum* were subjected to robust multichip average (RMA) analysis (65) and quantile normalization (63, 64) (Figure 3S). Transcription values for each gene were obtained using median polish summarization (54). Final transcription value for each sample was a mean of up to twelve technical replicates (Blocks 1 and 2 with up to six replicate probes in each block in the *T. erythraeum* microarray design). A gene was selected for further analysis if it had log_2_ transcription above 64 in at least 25% of samples and an interquartile range across all samples on the log_2_ scale of at least 0.5. This filtering resulted in 4128 genes, which were used in further analysis.

### 3) Comparison of diel transcription patterns for all cyanobacteria

Transcription data for *Prochlorococcus* sp. MED4, *Cyanothece* sp. ATCC 51142 and *Crocosphaera watsonii* WH 8501 was collected from previous published data (16–18). *Cyanothece* sp. ATCC 51142 and *C. watsonii* WH 8501 microarray data were downloaded from ArrayExpress (http://www.ebi.ac.uk/aerep/) using accession no. E-TABM-386 and E-TABM-737, respectively. The genes with periodic transcriptional patterns for all studied cyanobacteria (*Prochlorococcus* sp. MED4, *Cyanothece* sp. ATCC 51142, *C. watsonii* WH 8501, *T. erythraeum* and UCYN-A) were identified using the R package “cycle” based on Fourier analysis, and the genes with FDR<0.25 were selected for further comparison (66) (Table S2). To compare the diel transcription patterns among the cyanobacteria, gene transcription values for each cyanobacterium were selected for over 36 hours. Eight points were selected for UCYN-A (L6, L9, D3, D6, 2D12, 2L3, 2L9, 2L12), 9 points for *T. erythraeum* (D12, L3, L6, L9, L12, D3, D6, D9, 2D12), 6 points for *Cyanothece* sp. ATCC 51142 (L2, L6, L10, D2, D6, D10), 8 points for *C. watsonii* WH 8501 (D11, L1, L6, L11, D1, D6, 2D11, 2L1) and 19 points for *Prochlorococcus* sp. MED4 (D12 - 2L12 every 2 hours). L and D stand for light and dark period, respectively, 2L and 2D the second light-dark cycle, and the number the corresponding hours entering light or dark period. Because the studies had a few dissimilar sampling times, the missing values were interpolated using the Stineman algorithm implemented in the *imputeTS* package (67). A network was constructed based on the Pearson correlation and using ‘make_network’ function in phyloseq (68). The maximum distance between connecting nodes was selected as 0.5 unless otherwise noted in figure legends.

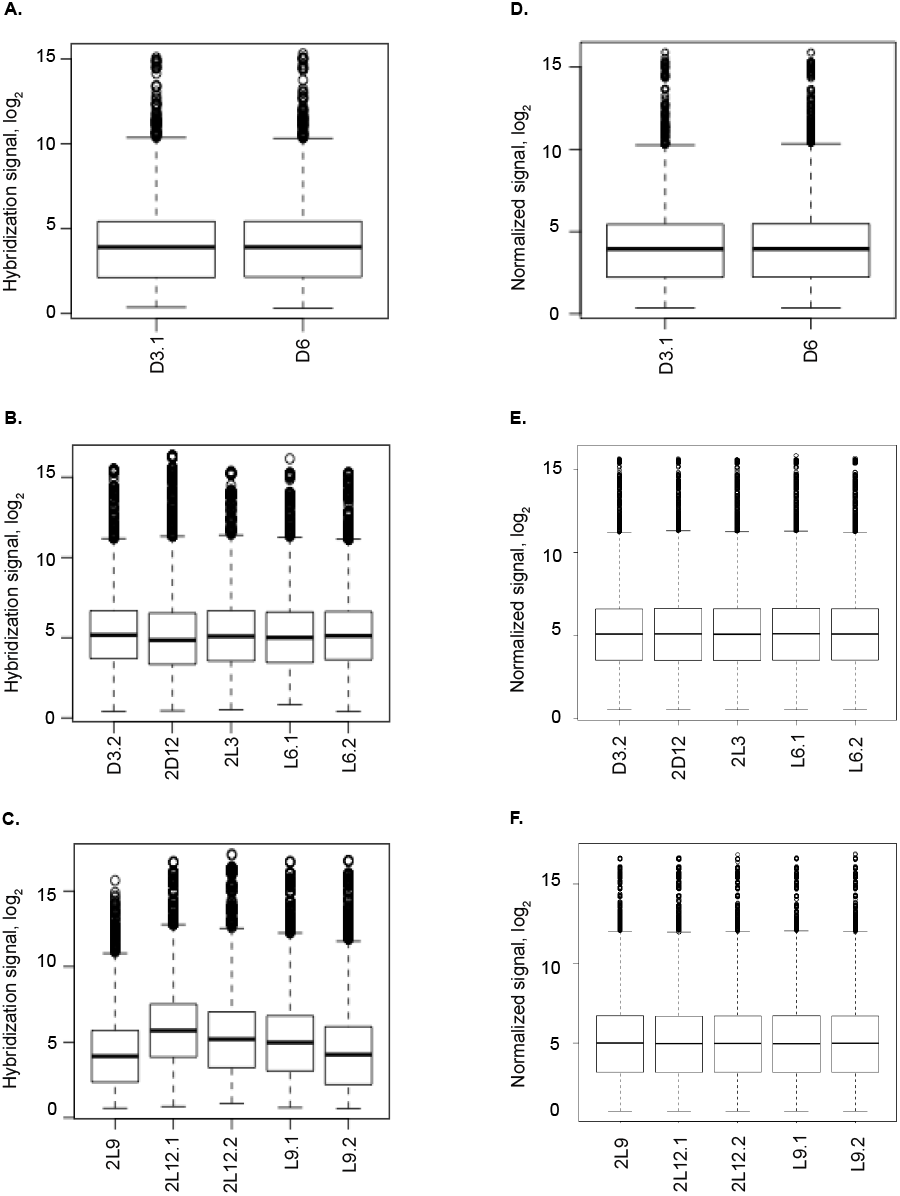

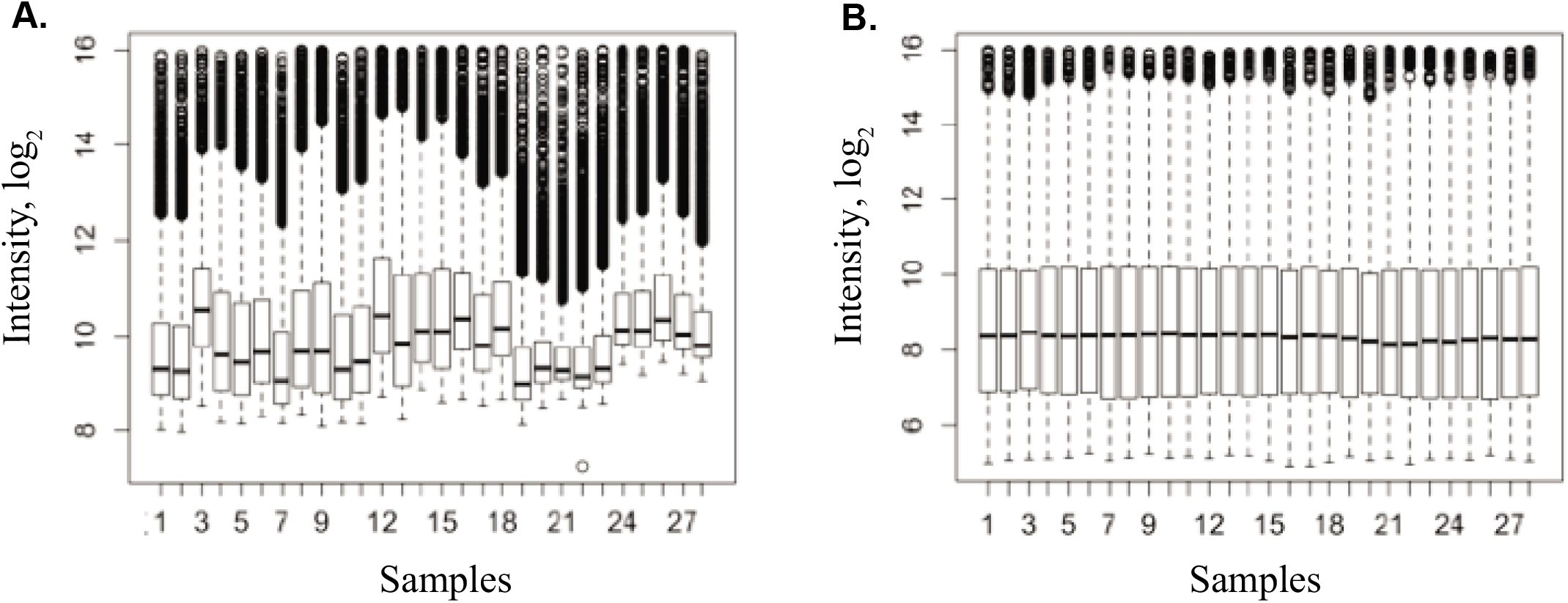

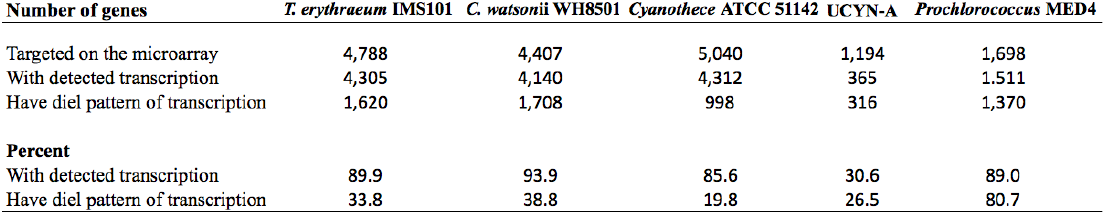

| **Number of genes** | *T. erythraeum* IMS101 | *C. watsonii* WH8501 | *Cyanothece* ATCC 51142 | UCYN-A | *Prochlorococcus* MED4 |
| --- | --- | --- | --- | --- | --- |
| Targeted on the microarray | 4,788 | 4,407 | 5,040 | 1,194 | 1,698 |
| With detected transcription | 4,305 | 4,140 | 4,312 | 365 | 1,511 |
| Have diel pattern of transcription | 1,620 | 1,708 | 998 | 316 | 1,370 |
| **Percent** |  |  |  |  |  |
| With detected transcription | 89.9 | 93.9 | 85.6 | 30.6 | 89.0 |
| Have diel pattern of transcription | 33.8 | 38.8 | 19.8 | 26.5 | 80.7 |

|  | Sampling<br>Time | Host associated<br>with 1 UCYN-A2<br>cluster<br>(cells ml<sup>-1</sup>) | Host associated<br>with 2 UCYN-A2<br>clusters<br>(cells ml<sup>-1</sup>) | Host associated<br>with 2 UCYN-A2<br>clusters<br>(%) |
| --- | --- | --- | --- | --- |
| *<u>Diel cycle 1</u>* | 12 pm | 36.5 | 0.0 | 0.0 |
|  | 3 pm | 38.0 | 0.0 | 0.0 |
|  | 6 pm | 9.1 | 0.0 | 0.0 |
|  | 9 pm | 12.2 | 15.2 | 55.6 |
|  | 12 am | 6.1 | 9.1 | 60.0 |
|  | 3 am | 41.0 | 10.6 | 20.6 |
|  | 6 am | 7.6 | 0.0 | 0.0 |
|  | 9 am | 38.0 | 0.0 | 0.0 |
| *<u>Diel cycle 2</u>* | 12 pm | 44.1 | 0.0 | 0.0 |
|  | 3 pm | 53.2 | 0.0 | 0.0 |
|  | 6 pm | 36.5 | 0.0 | 0.0 |
|  | 9 pm | 16.7 | 28.9 | 63.3 |
|  | 12 am | 34.9 | 51.7 | 59.6 |
|  | 3 am | 51.7 | 4.6 | 8.1 |
|  | 6 am | 66.8 | 0.0 | 0.0 |
|  | 9 am | 27.3 | 0.0 | 0.0 |

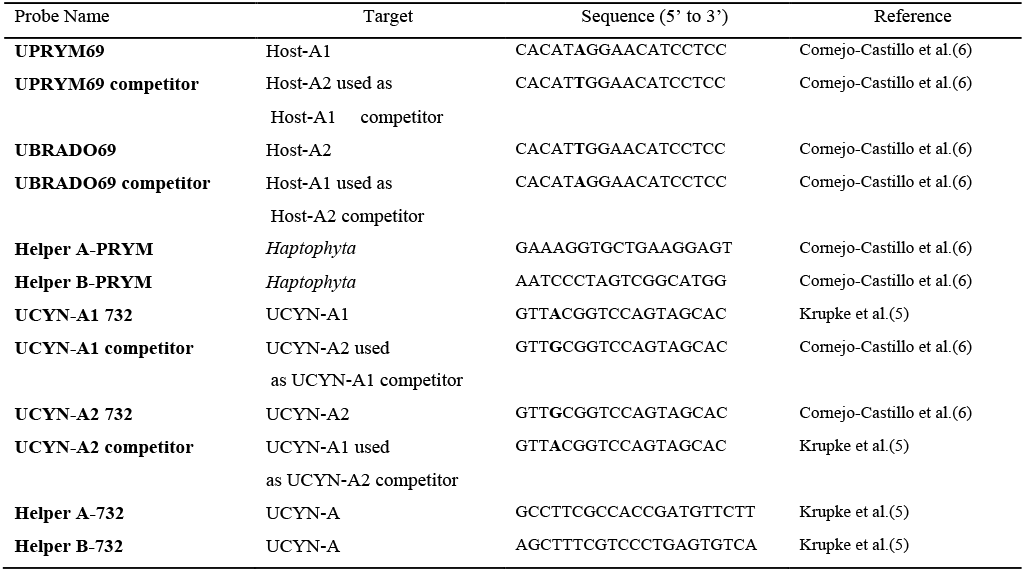

## Acknowledgments

We thank J.C. Meeks (University of California, Davis) for discussions, F. Azam (Scripps Institution of Oceanography, UC San Diego) for access to Scripps facilities, and K. Turk-Kubo and M. Hogan for lab and field support. J.M. García-Fernández and J. Díez-Dapena (University of Córdoba, Spain) and Marine Landa (University of Santa Cruz, CA, USA) for helping us to improve the manuscript. Microarray data have been deposited at NCBI Gene Expression Omnibus (GEO) under accession numbers GSE100124 and GSE99896. The following secure token has been created to allow review of record GSE100124 while it remains in private status: ufchawekhzmlpaj.

## Author contributions

M.M.M. designed the UCYN-A array, designed and performed the research and analyzed the data. I.N.S. analyzed the *T. erythraeum* array data, aided with the design of the UCYN-A array and comparison of transcription among cyanobacteria. T.S. designed the *T. erythraeum* array and performed the diel sampling of *T. erythraeum* cultures. H.F. aided sampling diel UCYN-A samples and performed the phylogenetic tree. A.M.C carried out and counted the CARD-FISH diel samples. J.P.Z. conceptualized the study, and M.M.M., I.N.S., T.S., H.F. and J.P.Z. drafted and edited the manuscript and figures. All authors read and approved the final manuscript.

